# The microbiota regulates inflammatory responses to toxin-induced CNS demyelination but has minimal impact on remyelination

**DOI:** 10.1101/575829

**Authors:** Christopher E McMurran, Alerie Guzman de la Fuente, Rosana Penalva, Ofra Ben Menachem-Zidon, Yvonne Dombrowski, Ginez A Gonzalez, Chao Zhao, Fynn N Krause, Adam MH Young, Julian L Griffin, Clare A Jones, Claire Hollins, Markus M Heimesaat, Denise C Fitzgerald, Robin JM Franklin

## Abstract

The microbiota is now recognised as a key influence on the host immune response in the central nervous system (CNS). As such, there has been some progress towards therapies that modulate the microbiota with the aim of limiting immune-mediated demyelination, as occurs in multiple sclerosis. However, remyelination – the regeneration of myelin sheaths – also depends upon an immune response, and the effects that such interventions might have on remyelination have not yet been explored. Here, we show that the inflammatory response during CNS remyelination in mice is modulated by antibiotic or probiotic treatment, as well as in germ-free mice. We also explore the effect of these changes on oligodendrocyte progenitor cell differentiation, which is inhibited by antibiotics but unaffected by our other interventions. These results reveal that high combined doses of oral antibiotics negatively influence remyelination and further our understanding of how mammalian regeneration relates to the microbiota.

## Introduction

Our knowledge of the microbiota and its relationship with host immunity, metabolism and neurobiology has significantly expanded in recent years (1). However, the role of the microbiota in mammalian regeneration remains relatively unexplored. As endogenous tissue regeneration is facilitated by a local immune response (2), it is feasible that intestinal microbes, which modulate the host immune system (3,4), could help shape the outcome of regeneration in a clinical setting. The gut microbial community can be easily modified using oral antibiotics or probiotics, thus addressing this question is of great clinical relevance.

Here, we investigate how the intestinal microbiota influences remyelination, the most effective form of regeneration observed in the mammalian central nervous system (CNS) and a promising therapeutic strategy for multiple sclerosis (MS) and other myelin diseases (5). Successful remyelination can bring functional recovery (6,7) and protect axons from degeneration (8). This success depends on an inflammatory response by microglia and infiltrating macrophages (9) that progresses and resolves appropriately (10,11). Due to its distance from any epithelial interface with environmental microbes, CNS remyelination is an ideal model to explore systemic influences of the microbiota on regeneration.

The rationale for our studies is based on the close relationship between remyelination and the innate immune response, and in turn between the immune system and the microbiota. To create an environment that permits remyelination, endogenous microglia and infiltrating macrophages must clear myelin debris remaining from the disintegrated sheath (12), and also secrete pro-regenerative factors (10,11,13). These roles hinge upon a coordinated immune response to demyelination, failure of which can impair remyelination, as occurs in older animals (14,15). Meanwhile, there is now a substantial body of evidence linking the microbiota to CNS inflammation across various contexts. For example, germ-free or antibiotics-treated mice have transcriptionally immature microglia, which have an impaired response to lipopolysaccharide (LPS) or viral stimulation (3), whilst antibiotic depletion of the microbiota reduces monocyte entry into the brain (4). The immune system is therefore a strong candidate for conveying an influence from the microbiota to regeneration in distant tissues such as the CNS.

There is a growing appreciation of the relationship between the microbiota and MS. Patients with MS have a distinct microbiome compared to healthy controls (16,17), and this may have a role in disease pathogenesis, given that faecal transplant from patients with MS can provoke immune-mediated demyelination in predisposed germ-free mice (18). Much focus has centred upon antibiotic or probiotic interventions that aim to limit demyelination in MS and animal models (19–22). However, the effects that such treatments might have on remyelination remain to be elucidated.

Here we use a variety of interventions to alter the murine microbiota, before quantifying inflammation and oligodendrocyte progenitor cell (OPC) activity in response to toxin-induced demyelination. Across all models, altering the microbiota was found to modulate the inflammatory response following demyelination, which was impaired in antibiotics-treated or germ-free mice and augmented in probiotic-treated mice. However, no clear relationship emerged between the microbiota, the OPC response to demyelination and subsequent remyelination.

## Results

### Combined antibiotics treatment to deplete the microbiota impairs inflammation following toxin-induced demyelination

To explore whether extensive microbial depletion would affect remyelination, C57BL/6 mice were administered broad-spectrum antibiotics (ABX) in their drinking water for 8 weeks (**Fig. 1A**). This combination of ampicillin/sulbactam, ciprofloxacin, vancomycin, metronidazole and imipenem has previously been shown to cause near-complete depletion of the microbiota (4), which we confirmed by quantitative RT-PCR of faecal DNA (**Fig. 1B**). Demyelination was then induced by focal injection of lysolecithin into the ventral white matter of the spinal cord.

**Figure 1:**
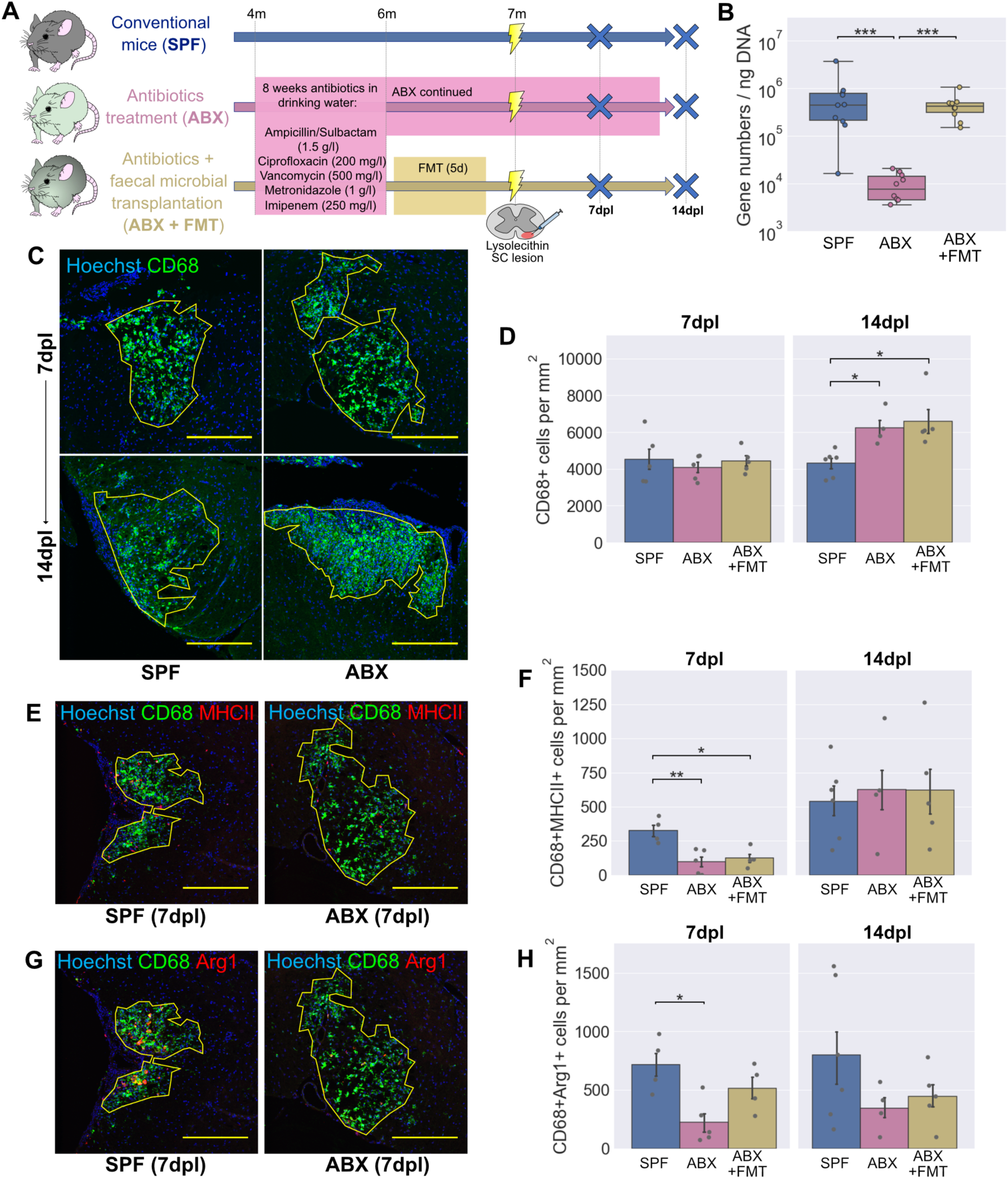
Antibiotics treatment to deplete the microbiota alters the inflammatory response following lysolecithin-mediated demyelination. (**A**) Schematic diagram of the experiment: mice were administered ABX in their drinking water for 2 months, after which one group received a faecal microbial transplant (FMT) whilst another group were continued on ABX. Mice were sacrificed at 7 and 14 days following lysolecithin injection into the ventral white matter of the spinal cord. (**B**) RT-PCR of faecal DNA showing substantial depletion of the microbiota by ABX and return to normal levels with FMT. (**C-D**) Representative images (**C**) and density (**D**) of CD68+ activated microglia / infiltrating macrophages within the lesion boundary (yellow line). (**E-F**) Representative images (**E**) and density (**F**) of MHCII+CD68+ activated microglia / infiltrating macrophages within lesions. (**G-H**) Representative images (**G**) and density (**H**) of Arg1+CD68+ activated microglia / infiltrating macrophages within lesions. Scale bars (**C**, **E**, **G**) = 250μm. Error bars show mean ± SEM; *p<0.05, **p<0.01, ***p<0.001; (**B**) Kruskal-Wallis with Dunn’s *post hoc* test; (**D**, **F**, **H**) one-way ANOVA with Tukey HSD *post hoc* test, n=4-6 mice.

To investigate baseline differences in microglia following ABX treatment, areas of unlesioned white matter were stained with an antibody to Iba1. Microglial morphology, which correlates with activity within a given tissue (3,23), was then analysed (**Fig. 1 supplement 1A**). Whilst a principal component analysis based on a number of morphological features showed clear separation between microglia from grey and white matter, there were no differences detected in microglia of unlesioned white matter following ABX treatment (**Fig. 1 supplement 1B**).

The inflammatory response within lesions was examined at 7 and 14 days post lesion (dpl) by staining for CD68 to label activated microglia and infiltrating macrophages. As subpopulations of microglia/macrophages have distinct effects in remyelination (11), we also quantified several markers expressed by subpopulations of these CD68+ cells. This panel consisted of two M1-associated markers (major histocompatibility complex class II (MHCII) and inducible nitric oxide synthase (iNOS)) and two M2-associated markers (arginase-1 (Arg1) and mannose receptor (MR)) (11,24).

At 7 dpl, lesions of ABX-treated mice were the same size as specific pathogen-free (SPF) controls (**Fig 1 supplement 1C**) and with the same total density of CD68+ microglia/macrophages (**Fig. 1C,D**). However, there were fewer CD68+ cells expressing either MHCII (**Fig. 1E,F**) or Arg1 (**Fig. 1G,H**). To determine the reversibility of this effect, ABX-treated mice received a faecal microbial transplant (FMT) from SPF mice by oral gavage to reconstitute their microbiota. With FMT treatment, Arg1 expression increased and was no longer significantly different from SPF control levels (**Fig. 1H**), whereas MHCII expression remained low (**Fig. 1F**). The other two other markers investigated (iNOS and MR) were not altered following ABX treatment (**Fig. 1 supplement 1D-G**). At a later timepoint, 14dpl, differences were no longer observed in either Arg1 or MHCII in the ABX-treated mice (**Fig. 1E-H**). However, the total number of CD68+ microglia/macrophages was increased following ABX treatment and not reversed by FMT (**Fig. 1C,D**).

Taken together, these results indicate that broad-spectrum antibiotic treatment results in dysregulated CNS inflammation following demyelination. Specifically, we observed an increase in total CD68+ cells from 7 to 14dpl in ABX-treated groups, not seen amongst SPF controls. Coupled with this, MHCII and Arg1 expression by microglia/macrophages in the ABX-treated groups were initially low, catching up to control levels with the increase in total CD68+ cells at 14dpl. Whilst microglia/macrophages expressing M1-or M2-associated markers are considered to have distinct functional effects during remyelination (11), we did not observe concomitant regulation of either M1-or M2-associated markers with our interventions, highlighting the intricacies of CNS inflammation in this context. Instead, our microbial depletion model was dominated by a delayed but aggressive pro-inflammatory innate immune response.

### Combined antibiotics treatment impairs myelin debris clearance and OPC differentiation

Central to CNS remyelination is the generation of new oligodendrocytes to restore the myelin sheath. These derive from endogenous OPCs, which migrate to lesions, proliferate and subsequently differentiate into oligodendrocytes (5,25). Defective differentiation from OPC to oligodendrocyte is commonly the bottleneck at which remyelination fails in human lesions and animal models (26–28). Microglia and infiltrating macrophages promote remyelination through the clearance of myelin debris, which blocks OPC differentiation in culture (29) and impairs remyelination in demyelinated lesions (12). We investigated whether the dysregulated inflammatory response following ABX treatment would be associated with reduced myelin debris clearance.

Lesions were stained with an antibody for a degraded epitope of myelin basic protein (dMBP), which becomes exposed in myelin debris (30) (**Fig. 2A,B**). At 7dpl, antibiotics did not affect the quantity of myelin debris in the lesion area. By 14dpl there had been an expected reduction in myelin debris in the SPF controls: however, this was less pronounced amongst the ABX-treated group, with more debris persisting at the later timepoint. The effect was reversed in the FMT group, which had a similar dMBP+ area to controls.

**Figure 2:**
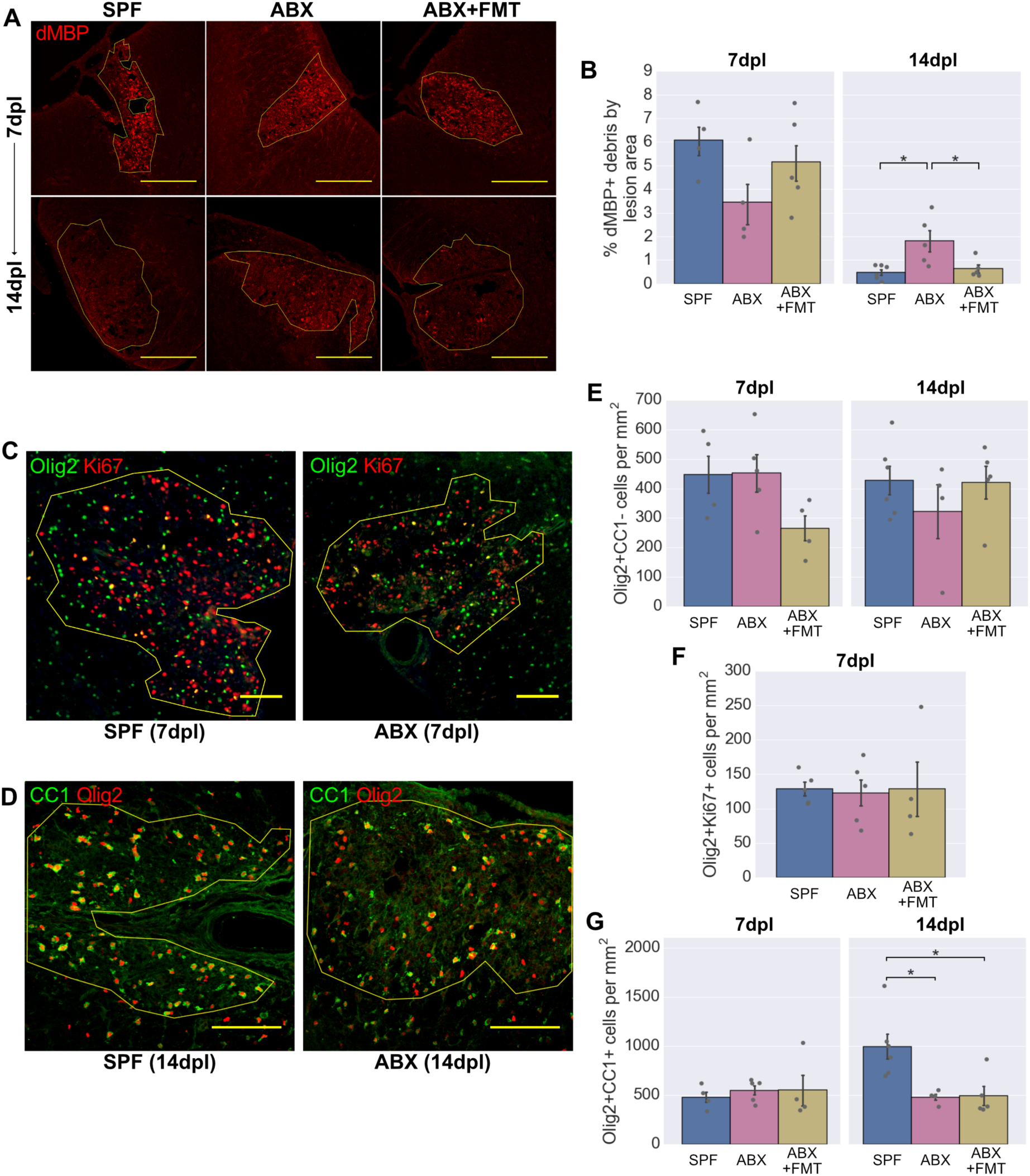
Fewer new oligodendrocytes are generated in the lesions of ABX-treated mice. (**A-B**) Representative images (**A**) and area (**B**) of dMBP+ myelin debris within lesions. (**C-D**) Representative images of Olig2+Ki67+ proliferating OPCs (**C**) and Olig2+CC1+ oligodendrocytes (**D**) within lesions. (**E-G**) Density of Olig2+CC1-OPCs (**E**), Olig2+Ki67+ proliferating OPCs (**F**) and Olig2+CC1+ oligodendrocytes (**G**) within lesions. Scale bars (**A**) = 250μm, (**C**, **D**) = 100μm. Error bars show mean ± SEM; *p<0.05; one-way ANOVA with Tukey HSD *post hoc* test, n=4-6 mice.

We next explored how the OPC response would be affected by ABX treatment, using the nuclear transcription factor Olig2 to label cells of the oligodendrocyte lineage and CC1 to distinguish differentiated oligodendrocytes (CC1+) from OPCs (CC1-). ABX did not affect the total number of OPCs present (**Fig. 2D,E**), nor the number of proliferating OPCs identified by Ki67 staining (**Fig. 2C,F**). However, there were fewer differentiated oligodendrocytes in the lesions of ABX-treated mice at 14dpl, suggesting a defect in OPC differentiation (**Fig. 2D,G**). This was not restored by FMT in the ABX-treated group.

In summary, following demyelination in ABX-treated mice, the altered inflammatory response is associated with impaired myelin debris clearance and impaired OPC differentiation. The effects of ABX treatment on remyelinating lesions could feasibly be caused by either ABX-mediated depletion of the microbiota or other off-target effects of this oral ABX regime. Two approaches were taken to explore these possibilities: 1) testing for off-target effects of ABX on relevant CNS cells in culture and 2) investigating remyelination in germ-free (GF) mice, which were reared in a sterile environment and thus were constitutionally devoid of microbes without ABX exposure.

### Combined antibiotics treatment does not affect microglial phagocytosis or OPC differentiation *in vitro*

Some antibiotics are able to penetrate and act directly within the CNS. One example is the broad-spectrum tetracycline, minocycline, which can directly inhibit microglia and influence remyelination (31–33). Four of the antibiotics we administered were considered to have sufficient bioavailability to reach the CNS in significant concentrations: ampicillin, ciprofloxacin, metronidazole and the β-lactamase inhibitor sulbactam. Available pharmacokinetic data (34–41) were used to estimate the concentrations these drugs would reach in the murine CNS (**Fig. 3 supplement 3A**), which were then applied to primary murine microglial cultures for 48 hours (**Fig. 3A**). Following ABX treatment, microglia were exposed to myelin debris and phagocytic uptake was quantified after 4 hours. None of the ABX individually, nor in combination, directly inhibited myelin phagocytosis by microglia (**Fig 3B,C**; **Fig 3 supplement 3B**).

**Figure 3:**
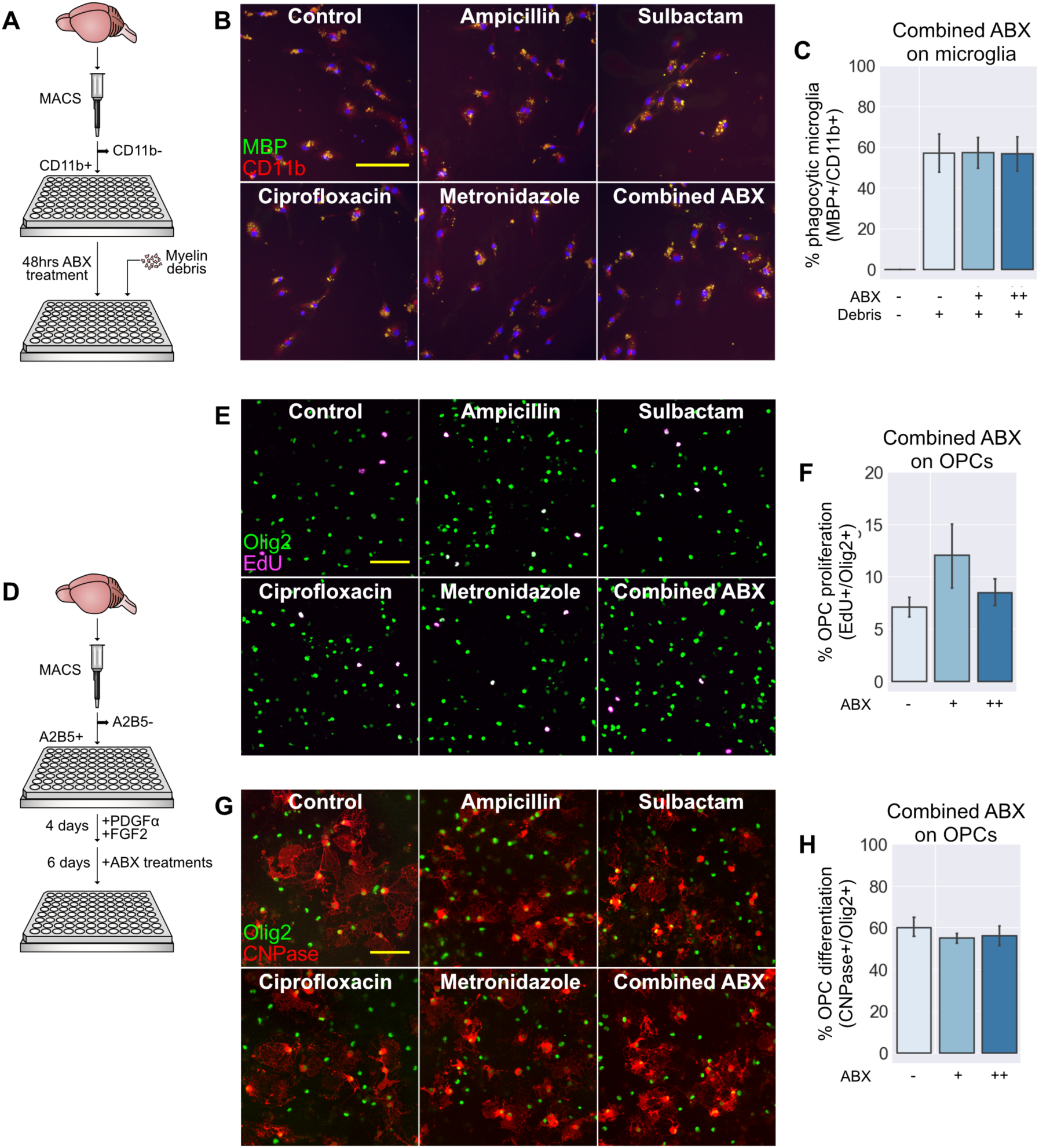
Exposure of microglia and OPCs to antibiotics *in vitro*. (**A**) Primary microglia were isolated from P6-8 mice using magnetic-activated cell sorting (MACS) for CD11b and cultured for 48 hours in the presence of antibiotics. Cells were fixed after exposure to myelin debris for 4 hours. (**B**) After fixation, cells were stained with an antibody to MBP to visualise myelin uptake by phagocytosis. (**C**) Myelin uptake following exposure to combined antibiotics at their estimated CNS concentration (++) or a 10% dose (+) was comparable to control levels. (**D**) Primary OPCs were isolated from P6-8 mice using MACS for A2B5 and cultured in the presence of growth factors. After 4 days, antibiotics were applied and growth factors were withdrawn to allow differentiation. Cells were fixed after 6 days of differentiation, following 3 hours of EdU exposure. (**E**) After fixation, cells were stained with an antibody to Olig2 and incorporated EdU was labelled to visualise OPC proliferation. (**F**) The doses of antibiotics used had no significant effect on OPC proliferation. (**G**) Cells were stained with antibodies to Olig2 and CNPase to visualise OPC differentiation. (**H**) The doses of antibiotics used had no significant effect on OPC differentiation. Scale bars (**B,E,G**) = 100μm. Error bars show mean ± SEM; paired-samples *t*-test with Holm-Bonferroni correction n=4-5 separate experiments.

It is also possible that ABX can act directly on OPCs to inhibit differentiation in lesions of mice receiving oral ABX. To test this, primary cultures of murine OPCs were treated with ABX and the effects on differentiation and proliferation were determined by immunocytochemistry after six days of differentiation conditions (**Fig. 3D-H**). ABX treatment had no effect on OPC proliferation as measured by EdU (**Fig. 3E-F**), nor was there a difference in expression of the differentiation marker CNPase (**Fig. 3G-H**). Similarly, applying each antibiotic alone had no effect on these parameters (**Fig 3 supplement 3C-D**).

### Germ-free mice have a deficient inflammatory response during remyelination

The results from cell culture systems suggest that the deficits observed in ABX-treated mice are not caused by direct effects of antibiotics on microglial or OPC functions critical for remyelination. However, antibiotics may have other indirect effects that are difficult to capture *in vitro*. To determine whether the microbiota can influence remyelination in the absence of ABX exposure, we investigated how remyelination would proceed in germ-free (GF) mice. As surgical lesions are impractical under the constraints of GF husbandry, we employed the cuprizone model, in which demyelination is induced by dietary administration of 0.2% cuprizone over a period of 5 weeks (**Fig. 4A**). The sterility of the GF mice was confirmed by faecal PCR for bacterial DNA (**Fig. 4B**) as well as routine in-house screening of animals and isolators using aerobic and anaerobic culture methods and microscopy of faecal smears. An ex-GF group were colonised with a microbiota after weaning, by co-housing with SPF controls, and this allowed us to distinguish developmental effects of lacking a microbiota from those reversible in adulthood. After cuprizone exposure, all mice were returned to a normal diet for a 3-week period of remyelination.

**Figure 4:**
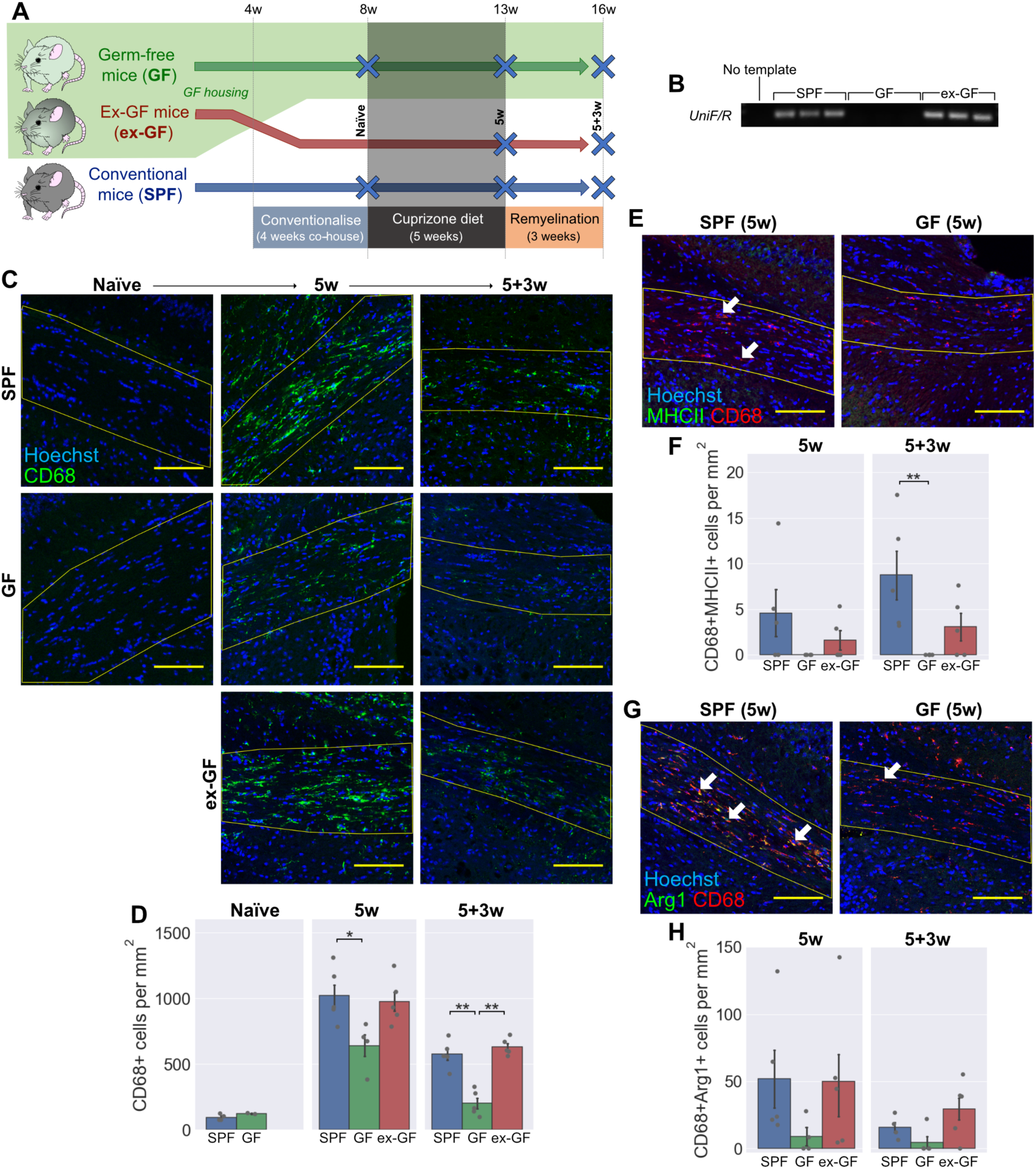
Germ-free mice have an altered inflammatory response following cuprizone-mediated demyelination. (**A**) Schematic diagram of the GF experiment: GF mice and SPF controls were fed a diet containing 0.2% cuprizone for 5 weeks from 8 weeks of age. Mice were sacrificed at the end of cuprizone administration, or after 3 weeks return to normal diet. A third group consisted of GF mice that were co-housed with SPF mice from 4 weeks of age, becoming colonised with a microbiota after weaning. (**B**) PCR for a universal prokaryotic 16S sequence in faecal DNA demonstrating absence of this amplicon in GF mice and its presence in the ex-GF group. (**C-D**) Representative images (**C**) and density (**D**) of CD68+ activated microglia / infiltrating macrophages within the corpus callosum (yellow line) in cuprizone naïve mice, following 5 weeks cuprizone exposure (5w) and following a further 3 weeks of normal diet (5+3w). (**E-F**) Representative images (**E**) and density (**F**) of MHCII+CD68+ activated microglia / infiltrating macrophages within the corpus callosum. (**G-H**) Representative images (**G**) and density (**H**) of Arg1+CD68+ activated microglia / infiltrating macrophages within the corpus callosum. Scale bars (**C**, **E**, **G**) = 100μm. Arrow heads (**E**, **G**) show representative double-positive cells. Error bars show mean ± SEM; *p<0.05, **p<0.01; (**D**) one-way ANOVA with Tukey HSD *post hoc* test, (**F**) Kruskal-Wallis with Dunn’s *post hoc* test; n=4-5 mice.

Like ABX-treated animals, GF mice had deficits in the inflammatory response that accompanied demyelination. The total number of CD68+ activated microglia and monocyte-derived macrophages was reduced in the GF group after the 5-week cuprizone treatment and remained so 3 weeks after cuprizone was withdrawn (**Fig. 4C,D**). Numbers of CD68+ cells were comparable to SPF controls in the ex-GF group, demonstrating that colonisation of GF mice can restore immune responsiveness and delineates an ongoing, dynamic role of the microbiota in the inflammatory response to tissue damage, rather than a critical window during development.

To investigate this inflammatory response in more detail, we stained for the same panel of microglia/macrophage markers as tested in the antibiotics experiment. Previously, ABX-treated mice were both found to have reduced expression of both MHCII and Arg1 during remyelination (**Fig. 1E-H**). Here, MHCII was similarly diminished in the remyelinating corpus callosum of GF mice, with negligible expression in microglia of any of these animals (**Fig. 4E,F**). There was no significant reduction in Arg1, despite a similar trend (**Fig. 4G,H**), nor was there a difference detected in iNOS or MR expression in GF mice (**Fig 4 supplement 4A-D**). A caveat for these negative results is that our study was powered to detect differences in OPC responses, and we observed considerable variability in the inflammatory responses between animals.

The deficits in the immune response of ABX-treated mice were associated with reduced myelin debris clearance after demyelination (**Fig. 2A,B**). A timecourse of cuprizone feeding in SPF mice showed that extensive dMBP+ myelin debris after 3 weeks of cuprizone diet is subsequently cleared by 5 weeks (**Fig 4 supplement 4E**). We subsequently compared dMBP staining at our 5w timepoint to determine whether myelin debris clearance was impaired under GF conditions. All groups showed similarly low levels of dMBP staining at 5w, indicating that any delay in debris clearance was not persistent beyond 5 weeks (**Fig 4 supplement 4E,F**).

In summary, we observed that the initial inflammatory response following demyelination in GF mice was altered in a similar pattern to ABX-treated animals, namely reduced numbers of CD68+ microglia/infiltrating macrophages with a corresponding reduction in MHCII expression. These changes point towards the microbiota being an important regulator of the immune response in the CNS during demyelination and remyelination. However, in our cuprizone model, the altered inflammatory response was not associated with a delay in myelin debris clearance after 5 weeks.

### Germ-free mice have normal OPC differentiation during remyelination

Next, the OPC responses during cuprizone-mediated demyelination and remyelination were examined in GF mice. Prior to cuprizone exposure, GF and SPF mice had comparable numbers of Olig2+CC1-OPCs (**Fig. 5A,B**) and Olig2+CC1+ oligodendrocytes (**Fig. 5A,C**) in the corpus callosum. Following 5 weeks cuprizone treatment, there was an 80% reduction in the density of Olig2+CC1+ oligodendrocytes in the SPF group (**Fig. 5C**). In contrast, the loss of oligodendrocytes after 5 weeks of cuprizone was less extensive amongst the GF group, whilst the ex-GF group were in line with SPF controls. At this timepoint, all groups experienced a similar rise in Olig2+CC1-OPCs (**Fig. 5B**), consistent with the phase of OPC migration and proliferation known to occur during the first 5 weeks of cuprizone administration (42). However, when we specifically quantified Olig2+Ki67+ proliferative OPCs at 5 weeks, this was higher in the GF mice than the other two groups (**Fig. 5D,E**).

**Figure 5:**
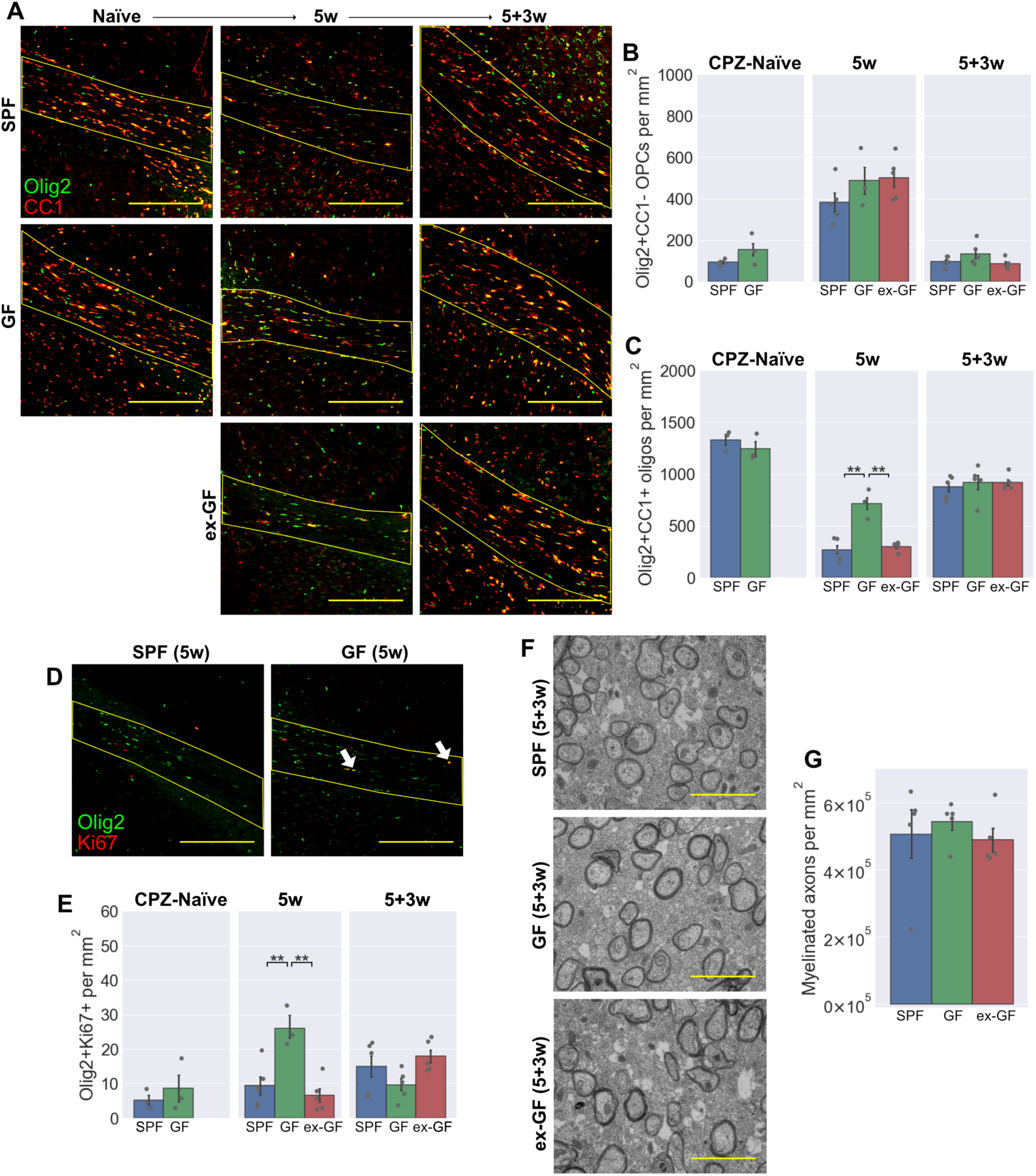
Germ-free mice have reduced oligodendrocyte loss following cuprizone administration, but no difference in OPC differentiation. (**A-C**) Representative images (**A**) and density of Olig2+CC1-OPCs (**B**) and Olig2+CC1+ mature oligodendrocytes (**C**) within the corpus callosum in cuprizone-naïve mice, following 5 weeks cuprizone exposure (5w) and following a further 3 weeks of normal diet (5+3w). (**D-E**) Representative images (**D**) and density (**E**) of Olig2+Ki67+ proliferating OPCs within the corpus callosum following 5 weeks cuprizone exposure. (**F-G**) Representative electron microscopy images (**F**) and quantification (**G**) of myelinated axons within the corpus callosum 3 weeks after cuprizone cessation. Scale bars: (**A,D**) = 200μm, (**F**) = 2μm. Arrow heads (**D**) show representative double-positive cells. Error bars show mean ± SEM; one-way ANOVA, n=3-5 mice.

Three weeks after cuprizone withdrawal, a timepoint at which remyelination should be progressing (42,43), all three groups exhibited a fall in OPC counts with a concurrent rise in differentiated oligodendrocytes. Importantly, there were no differences in OPC or oligodendrocyte numbers between the groups at this later timepoint (**Fig. 5B,C**), suggesting that there was no delay in OPC differentiation during the remyelination of GF mice.

Finally, the degree of remyelination following cuprizone cessation was investigated by transmission electron microscopy (**Fig. 5F**). The corpus callosum of SPF controls, GF and ex-GF mice all exhibited comparable densities of myelinated axons 3 weeks after cuprizone treatment was stopped, indicating that remyelination had progressed similarly between groups (**Fig. 5G**). The g-ratios of these axons were also measured and likewise revealed no differences (**Fig 5 supplement 5A,B**).

In summary, GF mice were partially resistant to the effects of cuprizone, with less extensive oligodendrocyte loss following 5 weeks of dietary administration. This effect could be related to their diminished innate immune response during cuprizone treatment (**Fig. 4**), given that various studies have demonstrated a role of the immune system, particularly microglia and neutrophils, in mediating cuprizone-induced demyelination (44,45). The late peak in OPC proliferation observed in the GF group (**Fig. 5D,E**), may similarly be in response to a sluggish innate immune response. However, 3 weeks after cuprizone was stopped, the absence of any difference in OPC or oligodendrocyte numbers combined with similar densities of myelinated axons suggest that remyelination itself is unaffected in GF mice.

### Probiotic VSL#3 can augment the innate immune response during CNS remyelination

Taking the results from the antibiotics and germ-free experiments, together with *in vitro* studies, an intact microbiota appears to be necessary for the appropriate inflammatory response during remyelination. However, the effect that microbial depletion has on OPC responses, and thus remyelination itself, remains less certain. We next explored the inverse of this relationship: whether the microbiota can be manipulated therapeutically to enhance appropriate inflammatory responses and subsequently remyelination. For this, we used older mice (aged 14 months) in which remyelination occurs more slowly with scope for improvement (14,46). For this study, mice were administered the probiotic VSL#3 by daily gavage for 1 month, prior to inducing focal demyelination in the spinal cord white matter by lysolecithin injection (**Fig. 6A**).

**Figure 6:**
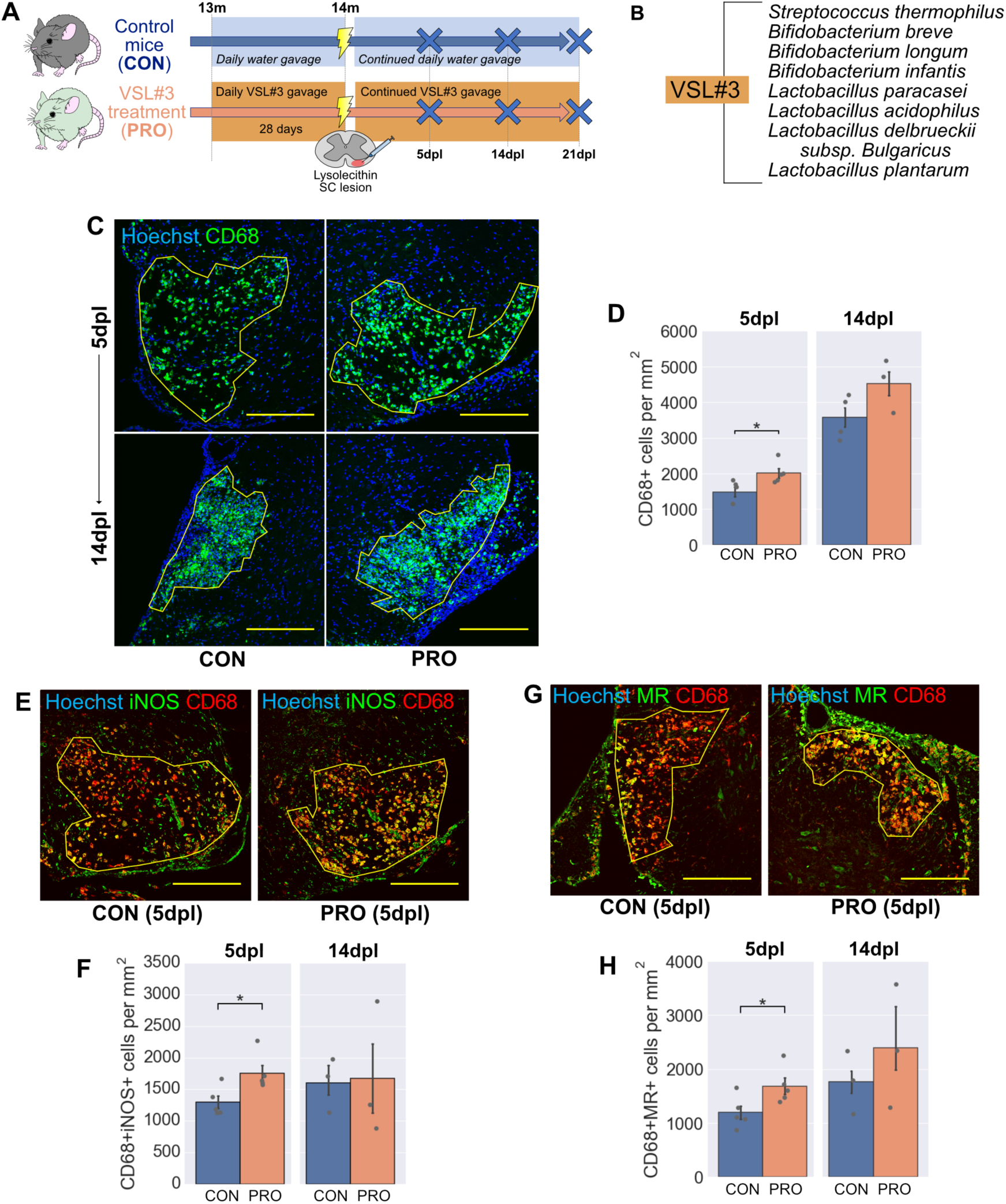
Probiotic VSL#3 enhances the onset of inflammation following demyelination. (**A**) Schematic diagram of the probiotic experiment: 13-month old mice were administered 1.35 x 109 colony-forming units (CFU) VSL#3 daily for 1 month by oral gavage. Mice were sacrificed at 5 and 14 days following lysolecithin injection into the ventral white matter of the spinal cord. (**B**) Constituent strains of VSL#3. (**C-D**) Representative images (**C**) and density (**D**) of CD68+ activated microglia / infiltrating macrophages within lesions. (**E-F**) Representative images (**E**) and density (**F**) of iNOS+CD68+ activated microglia / infiltrating macrophages within lesions. (**G-H**) Representative images (**G**) and density (**H**) of MR+CD68+ activated microglia / infiltrating macrophages within lesions. Scale bars (**C**, **E**, **G**) = 250μm. Error bars show mean ± SEM; *p<0.05; Student’s *t*-test, n=3-5 mice.

VSL#3, a freeze-dried formulation of 8 strains of Gram-positive bacteria (**Fig. 6B**), was chosen due to its good survival on transit through the GI tract (47) and several characterised therapeutic effects in the CNS (4,48,49). Additionally, VSL#3 enhanced concentrations of short chains fatty acids (SCFAs) in the faeces and serum of mice (**Fig. 6 supplement 6B,D**), consistent with a previous study (50). The SCFAs acetate, butyrate and propionate are microbial metabolites that are depleted in GF mice (**Fig. 6 supplement 6A,C**) and considered key signalling molecules in how the microbiota influence CNS inflammation (3,51).

Following VSL#3 treatment, there was a mild enhancement in the inflammatory response at 5dpl, with greater density of CD68+ activated microglia and infiltrating macrophages (**Fig. 6C,D**). When more specific inflammatory markers were quantified, both iNOS and MR were expressed by greater numbers of microglia/macrophages in lesions following probiotic treatment at 5dpl (**Fig. 6E-H**). No differences were detected at 14dpl, a timepoint at which debris should be largely cleared and newly differentiated oligodendrocytes initiate remyelination. This shows that although probiotics augmented the initial inflammatory response at 5dpl, this was not sustained, detrimental inflammation. It was noted that expression of the two markers initially reduced by ABX (MHCII and Arg1) were not significantly enhanced by probiotic treatment (**Fig. 6 supplement 6E-H**). However, there was a trend in both cases which could have been limited by sample size.

### VSL#3 does not improve the outcome of remyelination

Finally, we assessed whether this inflammatory enhancement during the early stages would contribute to a better outcome in remyelination. In contrast to ABX-treated mice, in which the reduced inflammatory response resulted in prolonged presence of myelin debris, myelin debris was not cleared any faster in the lesions of probiotic-treated mice (**Fig. 7A,B**). Consistent with this, there was no difference in the OPC response, with OPC number (**Fig. 7C,D**), proliferation (**Fig. 7 supplement 7A-B**) and differentiation of new oligodendrocytes at 14dpl (**Fig. 7C,E**) unchanged by probiotic treatment. There was a small increase in the number of oligodendrocytes present at 5dpl (**Fig. 7C,E**). As this timepoint is considered too early for significant OPC differentiation to occur, even in young adults (52) and taking to account that no difference was observed at 14dpl, the peak time for OPC differentiation, this likely reflects a reduction in oligodendrocyte death, rather than enhanced OPC differentiation.

**Figure 7:**
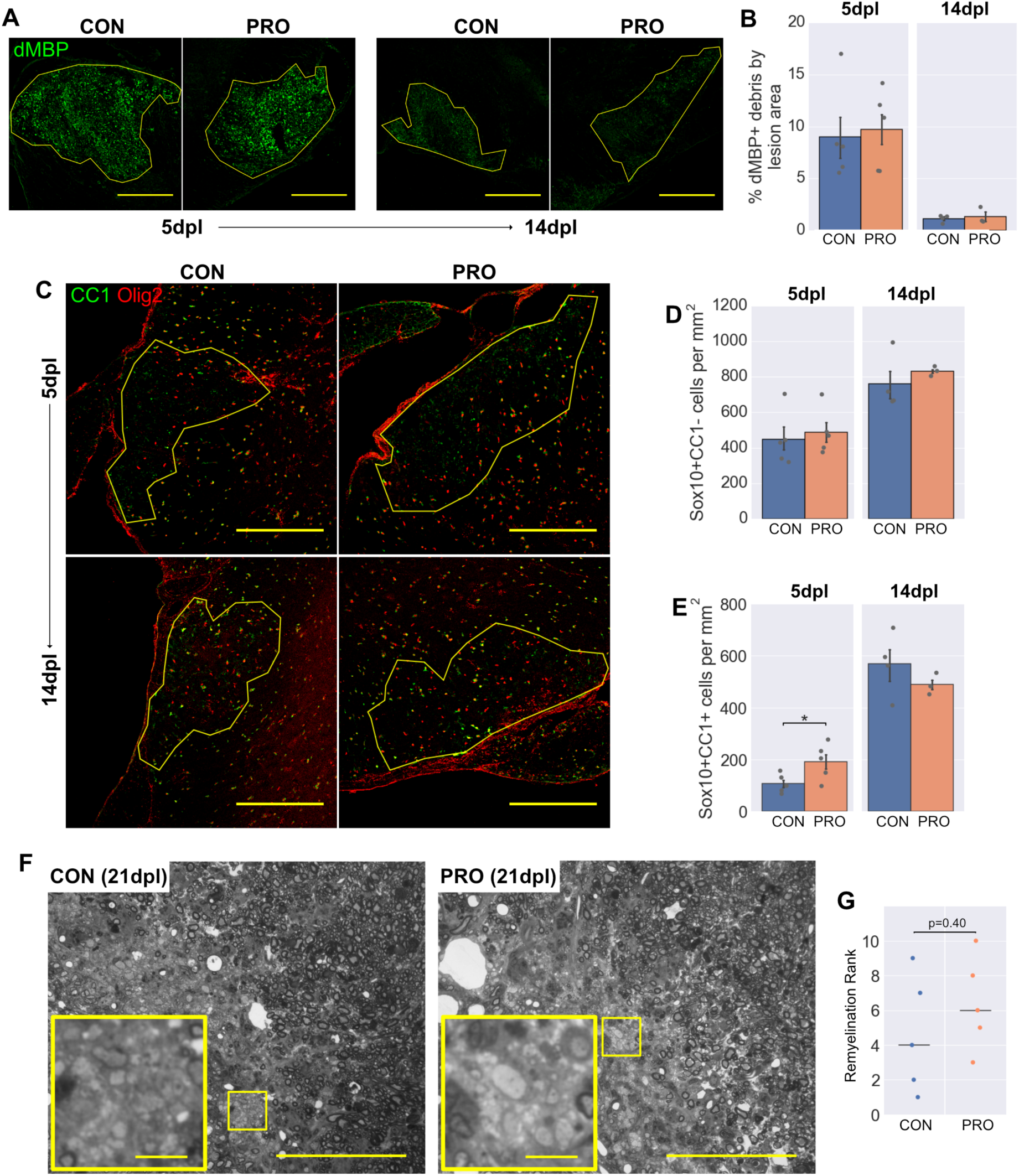
VSL#3 probiotic does not enhance remyelination in aged mice. (**A**) Representative images (**A**) and area (**B**) of dMBP+ myelin debris within lesions. (**C-E**) Representative images (**C**) and density of Sox10+CC1-OPCs and (**D**) and Sox10+CC1+ mature oligodendrocytes (**E**) within lesions. (**F**) Representative images showing the extent of remyelination assessed by toluidine blue staining of semi-thin resin sections. Insets are a 4x digital zoom of the boxed regions demonstrating persistent demyelination, typical of aged mice. Images are shown in grayscale. (**G**) Remyelination ranks assigned by a blinded assessor, with horizontal lines showing the median for each group. Scale bars (**A**, **C**) = 250μm, (**F**) = 100μm [inset scale bar = 10μm]. Error bars show mean ± SEM; (**B**, **D**, **E**) Student’s *t*-test, n=3-5 mice; (**H**) Mann–Whitney *U* test, n=5 mice.

To confirm that there was no therapeutic effect of VSL#3 on remyelination, semi-thin resin sections were taken at a later timepoint (21dpl) and myelin stained with toluidine-blue (**Fig. 7F**). The completion of remyelination was ranked by two blind assessors, neither of whom detected a difference in remyelination between groups (**Fig. 7G**).

Thus, administration of the probiotic VSL#3 promoted a stronger initial inflammatory response in aged mice following demyelination. However, this did not lead to faster clearance of myelin debris or OPC differentiation, and overall remyelination was unchanged compared to control mice.

## Discussion

The aim of these experiments was to explore how the microbiota can influence remyelination in the mammalian CNS. Across all three models, the responses of microglia and infiltrating macrophages were modulated by interventions that altered the microbiota. Broadly, this amounted to a blunting of the early immune response in ABX-treated or GF mice and an enhancement in the onset of inflammation when treated with the probiotic VSL#3. Alongside this, ABX-treated mice had deficits in myelin debris clearance and OPC differentiation. However, these findings were neither replicated in GF mice, nor reversed by probiotic administration. *In vitro*, ABX did not directly influence microglial phagocytosis or OPC proliferation and differentiation, though other indirect effects of ABX are difficult to exclude. Bringing together these findings from different models, the microbiota consistently shapes the inflammatory response during CNS remyelination, but the effect of this on regenerative responses by OPCs is limited.

Interactions between microbiota and regeneration have been observed in other tissues where contact is more direct. Skin wounds heal faster in GF mice, associated with reduced neutrophil accumulation (53), whilst microbiota-derived lipopolysaccharide (LPS) was found to dictate regenerative responses in the intestinal crypt (54). Similarly, the regenerative capacity of planaria was diminished by pathogenic shifts in their microbiota (55). However, it seems likely that such interactions become less relevant at anatomically distant sites – for example, the evidence linking liver regeneration to the microbiota is more contentious (56,57). Outside a regenerative context, changes in developmental or adaptive myelination in the CNS have previously been linked to the microbiota. GF mice are hypermyelinated specifically in the prefrontal cortex (PFC) (58), whilst a PFC *hypo*myelination phenotype could be transferred between different strains of mice by FMT (59). Whilst a role for the microbiota in myelin homeostasis is exciting, the findings of both studies were region-specific and occur in a very different context to remyelination, which is accompanied by tissue damage and a robust inflammatory response that directly influences the efficiency of regeneration.

Though neither our GF nor probiotic study demonstrated a difference in remyelination, in both cases there was a protective effect of the intervention on demyelination, as seen by increased numbers of oligodendrocytes at timepoints too early for substantial OPC differentiation to have occurred. This could be explained by the role of the immune system in mediating demyelination as well as remyelination in the lysolecithin and cuprizone models (44,45,60,61). Similarly, GF mice were resistant to immune-mediated CNS pathology in a model of Parkinson’s disease (51), whilst changes in the immune response with VSL#3 treatment were associated with white matter preservation in a spinal cord contusion model (49). As the immune system plays a prominent role in demyelination in MS, the majority of previous work on the gut-brain axis has focused on interventions that can limit inflammation and demyelination. For example, in the immune-driven experimental autoimmune encephalomyelitis (EAE) model, GF or antibiotics-treated mice are resistant to demyelination (19,62,63).

In conclusion, we have highlighted the microbiota as a key determinant of the inflammatory response during CNS remyelination. However, the regenerative responses of OPCs were largely independent of our interventions. The exception to this was following broad-spectrum oral antibiotic treatment, during which antibiotics do not inhibit OPCs directly, but may have other off-target systemic effects. These findings identify high, combined doses of oral antibiotics as a negative influence on remyelination and further our understanding of the interaction between microbiota and mammalian regeneration.

## Materials and methods

### Animal work

All animal work complied with the requirements and regulations of the United Kingdom Home Office (Project Licences: 70/7715 and 2789) or the European Guidelines for animal welfare with approval from the “Landesamt für Gesundheit und Soziales” (LAGeSo, Berlin registration number: G0184/12). Each study also adhered to the respective institutional guidelines, with approval by the local Animal Welfare and Ethical Review Body (AWERB). The antibiotics study was conducted at Queen’s University Belfast and Charité – Universitätsmedizin Berlin, the germ-free study at the University of East Anglia and the probiotic study at the University of Cambridge. Three to five mice of the same group were housed together in a cage, with *ad libitum* access to water and standard chow diet (other than during cuprizone administration) in a 12 hour light/dark cycle.

#### Focal lysolecithin lesions in antibiotics-treated mice

A cocktail of antibiotics was administered via the drinking water to 4-month old female C57BL/6 mice for 8 weeks. The antibiotics used were: ampicillin/sulbactam (1.5g/L), ciprofloxacin (200mg/L), vancomycin (500mg/L), metronidazole (1g/L) and imipenem (250mg/L) – a regime previously employed to cause virtually complete depletion of microbes in the gut (4,64). The faecal transplant group were then orally gavaged with faecal material from control mice, once per day for 5 days, whilst the antibiotics group continued to receive the antibiotics for the duration of the experiment. The control group were housed in specific pathogen free (SPF) conditions throughout. Mice were randomly allocated between these three groups at the start of the experiment. At age 7 months, demyelination was initiated by stereotactic injection of 1μl lysolecithin (L4129, Sigma-Aldrich, UK) into the thoracic spinal cord ventral white matter, under isoflurane anaesthesia. At 7 or 14 dpl, mice were intracardially perfused with 4% paraformaldehyde (PFA), the spinal cord was then dissected, post-fixed in cold PFA for 4-6 hours before overnight cryoprotection with 20% sucrose and OCT embedding. Tissue was cut on a cryostat in 12µm thick sections.

#### Cuprizone administration to germ-free mice

Male C57BL/6 mice were bred from a GF nucleus colony at the University of East Anglia and maintained in a flexible-film isolator, supplied with sterilised air, food, water and bedding. The GF group were maintained in these conditions throughout the experiment and compared to aged-matched SPF controls. The ex-GF mice were littermates of the GF group and remained in GF conditions until after weaning (4 weeks old), at which point they acquired a microbiome by co-housing with the SPF mice. Animals and isolators were monitored by routine in-house screening using aerobic and anaerobic culture methods, as well as microscopy of faecal smears. Demyelination was initiated by administration of a 0.2% cuprizone diet (TD.140803, Envigo, Huntingdon, UK), which received 50kGy γ-irradiation for sterilisation. All groups received the cuprizone diet for 5 weeks beginning at age 2 months in place of their regular diet and were then returned to regular diet for 3 weeks afterwards. Cuprizone diet was stored sealed at 4°C and replaced every 2-3 days during administration. During the 5-weeks of cuprizone diet, mice were regularly weighed and if substantial weight loss occurred (>15%), the diet was supplemented with small quantities of normal chow, given equally across all three groups. Mice were perfused with in the same manner as the antibiotics study either before the initiation of cuprizone (naïve), after 5 weeks cuprizone diet (5w) or after a subsequent 3 weeks of normal diet (5+3w). The brain was retrieved and bisected sagittally, with the left hemisphere post-fixed in PFA for immunohistochemistry (IHC) and the right hemisphere post-fixed in 4% glutaraldehyde for electron microscopy analysis. For IHC, 20μm coronal sections of the left hemisphere were taken from bregma −1 to 1mm.

#### Focal lysolecithin lesions in probiotic-treated mice

13-month old female C57BL/6 mice were administered a daily dose of 1.35×10^9^ colony forming units of VSL#3 (a gift from Janine DeBeer, Ferring Pharmaceuticals, London, UK), suspended in 100μl autoclaved water by oral gavage for 28 days. These were compared to age-matched control mice, which were instead gavaged daily with 100μl water. Randomly allocated control and probiotic-treated mice were housed in separate racks necessitating a change of gloves between handling, to avoid cross-contamination. Following this treatment, all groups received a focal injection of lysolecithin as described in the antibiotics study. Gavage treatments were recommenced from two days after this surgery and mice were sacrificed by perfusion-fixation at 5 or 14 dpl with 4% PFA or at 21 dpl with 4% glutaraldehyde.

### Immunohistochemistry

Cryosections on glass slides were brought to room temperature (RT) and rehydrated with PBS, prior to an antigen retrieval step, consisting of a 10-minute RT incubation with boiling antigen retrieval solution (Dako, Ely, UK). Slides were washed three times with PBS and blocked for 1 hour with 5% Normal Donkey Serum (NDS, Sigma-Aldrich) and 0.3% triton in PBS. Sections were then incubated with primary antibodies diluted in blocking solution in a humidity chamber overnight at 4°C. Primary antibodies: goat anti-Arg1 1:200 (sc-18351, Santa Cruz, Dallas, TX), mouse anti-CC1 1:100 (OP80, Calbiochem, San Diego, CA), rabbit anti-CD68 1:500 (ab125212, Abcam, Cambridge, UK), rat anti-CD68 1:200 (MCA1957, Serotec, Kiddlington, UK), rabbit anti-dMBP 1:500 (AB5864, Millipore, Burlington, MA), rabbit anti-Iba1 1:500 (019-10741, Wako, Osaka, Japan), rabbit anti-iNOS 1:200 (ab136918, Abcam), rabbit anti-Ki67 1:300 (ab15580, Abcam), rat anti-MHC class II 1:500 (14-5321-82, eBioscience, Thermo Fisher Scientific, San Diego, CA), goat anti-mannose receptor 1:500 (AF2535, R&D Systems, Minneapolis, MN), rabbit anti-Olig2 1:500 (AB9610, Millipore), goat anti-Sox10 1:200 (sc-17342, Santa Cruz). Slides were washed three times with PBS and incubated with relevant fluorophore-conjugated secondary antibodies (all Invitrogen, Thermo Fisher Scientific) diluted 1:500 in blocking solution for 2 hours at RT and protected from light. Secondary antibodies: Alexa 488 donkey anti-mouse (A21202), Alexa 488 donkey anti-rabbit (A21206), Alexa 488 donkey anti-rat (A21208), Alexa 568 donkey anti-goat (A11057), Alexa 568 donkey anti-mouse (A10037), Alexa 568 donkey anti-rabbit (A10042), Alexa 647 donkey anti-mouse (A31571), Alexa 647 donkey anti-rabbit (A31573). Nucleic acids were stained by 10 minutes incubation with 1μg/ml Hoechst. After three more PBS washes, sections were washed with water then mounted using Fluoromount-G (Southern Biotech, Birmingham, AL) with glass coverslips (VWR, Lutterworth, UK). Where mouse-derived primary antibodies were to be used on mouse tissue, a mouse-on-mouse blocking kit (Vektor, Peterborough, UK), was used according the manufacturer’s instructions.

Fluorescently immunolabelled sections were imaged using a Leica-SP5 confocal microscope with LAS software (Leica Microsystems, Wetzlar, Germany). Depending on the staining, either Z-stacks spanning the section widths were obtained and combined, or a single planar image was taken, and either a 20x or 40x lens was used.

### Histological analysis of remyelination

To produce semi-thin resin sections for toluidine blue staining, glutaraldehyde-fixed tissue was dissected into pieces of maximum 1 mm thickness and stained with 2% osmium tetroxide overnight at 4°C. Samples were then processed into resin (TAAB Laboratories Equipment Ltd., Aldermaston, UK), after which 0.75μm sections were cut using a microtome (Leica RM 2065) and stained with 1% toluidine blue.

These samples were further analysed by electron microscopy. 0.75μm sections were stained with aqueous 4% uranylacetate and lead citrate and visalised on a Tecnai G2 80-200keV transmission electron microscope. A minimum of five micrographs per animal were captured at 5000x from the splenium of the corpus callosum, close to the midline.

### Cell culture

#### Microglia isolation and culture

Microglia were isolated from P6-8 C57BL/6 mouse pups using a Magnetic-Activated Cell Sorting (MACS) protocol (**Fig. 3A**). Pups were euthanised by overdose of Pentoject (Animalcare Ltd., Hull, UK) and brains from 2-3 pups were pooled per replicate. Tissue was diced into small pieces and incubated at 37°C for 30 minutes in a dissociation solution, consisting of 34 U/ml papain (Worthington, Lakewood, NJ) and 20 μg/ml DNAse (Gibco, Thermo Fisher Scientific) in HALF (Hibernate-A equivalent, made in house). After this, tissue was triturated with a fire-polished glass pipette, passed through a 70μm cell strainer (Millipore) and centrifuged for 20 minutes at 800g in 22.5% Percoll (GE Healthcare, Little Chalfont, UK). The pellet, containing single cells, was labelled with magnetic bead-conjugated antibodies for CD11b (Miltenyi Biotech, Woking, UK), and CD11b+ microglia were eluted by MACS according the manufacturer’s instructions. Microglia were cultured at 10^4^ cells per well of a poly-D-lysine-coated 96-well microplate (Corning, NY), in DMEM/F12 (Gibco) supplemented with 10% foetal bovine serum (FBS, Biosera, Heathfield, UK), 2% B27, 500μM N-acetylcysteine and 1% penicillin-streptomycin. After 48 hours, media was changed to macrophage serum-free medium (Thermo Fisher Scientific), containing the antibiotic treatments. Following a further 48 hours, 10μg/ml myelin debris was added to each well for 4 hours. Myelin debris had been isolated from 2-3 month old C57BL/6 mice by discontinuous sucrose gradient centrifugation, as previously described (12,15). After this incubation, un-internalised debris was removed by washing with cold PBS and cells were fixed with 4% PFA.

#### OPC isolation and culture

OPCs were isolated from P6-8 C57BL/6 mouse pups using a MACS protocol (**Fig. 3D**), which was identical to the microglia protocol until the labelling stage. Cells were instead incubated with 1:1000 anti-A2B5 (Millipore) and then a magnetic bead-coupled goat-anti-mouse-IgM antibody (Miltenyi Biotech), prior to MACS. A2B5+ OPCs were cultured at 5000 cells per well of a poly-D-lysine and laminin-coated 96-well microplate, in DMEM/F12 (Gibco) supplemented with 5μg/ml insulin (Gibco), 1x Trace Elements B (Corning), 100μg/ml apo-transferrin, 16.1μg/ml putrescine, 40ng/ml sodium selenite, 60ng/ml progesterone, 60μg/ml N-acetylcysteine and 1% penicillin-streptomycin, 5μM forskolin and 1ng/ml biotin (all Sigma-Aldrich unless otherwise indicated). Cultures received growth factors for the first 4 days (10ng/ml/day of platelet-derived growth factor α (PDGFα) and fibroblast growth factor 2 (FGF2)), after which antibiotic treatments were introduced for a further six days (replaced after 3 days) in the absence of growth factors to allow differentiation. Cells were then incubated with 10μM 5-ethynyl-2’-deoxyuridine (EdU) for 3 hours prior to fixation with 4% PFA.

#### Antibiotic treatments *in vitro*

To apply antibiotic treatments (all Sigma-Aldrich) at doses approximating those of *in vivo* exposure, steady state plasma concentrations (*C*^*SS*(*P*)^) were estimated for each antibiotic administered to mice in their drinking water (**Fig. 3 supplement 3A**). These estimations were based on literature values of oral bioavailability (*F*), clearance (*CL*), and area-under-the-curve ratio of cerebrospinal fluid (CSF) to plasma (*AUC*_*CSF*_/*AUC*_*P*_), for ampicillin and sulbactam (34–36), ciprofloxacin (37–39) and metronidazole (39–41). Daily water consumption was assumed to be 4 ml/day/mouse.

#### Immunocytochemistry

Fixed cells in 96-well plates were blocked for 1 hour with 5% NDS and 0.1% triton in PBS. For the cell proliferation assay, EdU was labelled at this point using the Click-iT EdU Alexa Fluor 647 Imaging Kit (Thermo Fisher Scientific). Cells were then incubated with primary antibodies diluted in blocking solution overnight at 4°C. Primary antibodies: mouse anti-CD11b 1:300 (MCA275R, Serotec), mouse anti-CNPase 1:500 (C5922, Sigma-Aldrich), rat anti-MBP 1:500 (MCA409S, Serotec), rabbit anti-Olig2 1:500 (AB9610, Millipore). Cells were then washed three times with PBS and incubated with relevant fluorophore-conjugated secondary antibodies (see “Immunohistochemistry”) diluted in blocking solution for 1 hour at RT and protected from light. Nucleic acids were stained by 10 minutes incubation with 1μg/ml Hoechst, followed by three more PBS washes. Images were acquired automatically using an InCell2200 (GE Healthcare, Chicago, IL) or a Cell Insight CX5 (Thermo Fisher Scientific).

### Faecal polymerase chain reaction (PCR)

To determine the microbial load of antibiotics-treated mice, DNA from faecal pellets was extracted as described previously (4), quantified using QuantiT PicoGreen reagent (Invitrogen) and adjusted to 1 ng/μl. Copy numbers of the 16S rRNA gene were quantified by quantitative RT-PCR using generic eubacterial primers (Tib MolBiol, Berlin, Germany), and expressed per ng of total DNA.

To confirm microbial status of GF and ex-GF mice, DNA from faecal pellets was isolated using a QIAamp DNA Stool Mini Kit (Qiagen, Hilden, Germany) following the manufacturer’s instructions. 1μl eluted DNA was added to 24μl PCR SuperMix (Thermo Fisher Scientific) with 200nM *UniF/R* primer pairs (65), which recognise a 147bp conserved region of bacterial 16S rRNA. The conditions for the PCR were 95°C for 5 minutes, then 25 cycles of 95°C for 30s, 52°C for 30s and 72°C for 45s, and finally 72°C for 7 minutes. The PCR products were then separated on a 1% agarose gel and visualised under UV light.

### Detection of faecal / serum metabolites using GC-MS

Fresh faecal pellets were collected and stored in sterile Eppendorf tubes at −80°C. Serum samples were obtained from 100-200μl left ventricular blood collected prior to perfusion. The blood samples were incubated in sterile Eppendorf tubes for 1 hour at RT to allow coagulation, then separated by centrifugation for 15 minutes at 1500g. Serum was collected and stored at −80°C.

Short chain fatty acids (SCFAs) were extracted using a modified Bligh and Dyer method (66). In short, 15-25mg of faeces or 20µl of serum was transferred to a pre-chilled plastic tube and extracted with ice-cold 2:2:1 methanol:chloroform:water containing internal standard, following vigorous mixing, sonication and centrifugation (16,000g, 20 minutes). For serum samples, 100µl of the aqueous phase was transferred to pre-chilled glass tubes. Faecal samples were extracted twice, and a combined total of 300µl aqueous phase was transferred to pre-chilled glass tubes. Aqueous phases were dried down under nitrogen at 4°C, and derivatised as described previously (67).

SCFAs were measured by gas chromatography-mass spectrometry (GC-MS), on a Trace GC Ultra coupled to a Trace DSQ II mass spectrometer (Thermo Fisher Scientific). Derivatised samples were diluted 1:1 with hexane, and 2µl was injected onto a 50m x 0.25mm (5% phenyl-arylene, 95% dimethylsiloxane) column with a 0.25µm ZB-5MS stationary phase (Phenomenex, Macclesfield, UK). Full-scan spectra were collected at three scans per second over a range of 50 to 650 *m/z*. Data processing was carried out using *Xcalibur* (version 2.2, Thermo Fisher Scientific), and peaks were assigned based on the *National Institute of Standards and Technology* (USA) library. All solvents were of HPLC-grade or higher.

### Image analysis

Cell counts were semi-automated, using a combination of *Fiji, CellProfiler* and *CellProfiler Analyst* software (68). *Fiji* was used to create maximum projections of Z-stacks acquired by confocal microscopy. The region of interest (ROI i.e. the lesion area) was manually defined and individual channels extracted. These images were imported into *CellProfiler*, where the nuclear channel (Hoechst) was cropped to the pre-defined ROI, and nuclei were identified as primary objects. Other channels were normalised to the median background intensity to correct for variability in staining. A number of intensity features were measured for the nuclear and peri-nuclear regions of each cell in every channel, and this data was exported as a master database file. In *CellProfiler Analyst*, a training set of >50 cells was specified per group and used to train a classifier based on the feature set. The training data was increased until the classifier gave consistently comparable results to manual counting in sample images. Finally, “per lesion” cell counts were extracted for each image using this classifier.

To quantify the area of a lesion occupied by myelin debris, images of tissue stained for dMBP were imported into a *CellProfiler* pipeline, which applied a threshold to each image determined by the background (median) staining. The area of the image above this threshold was considered positive for myelin debris and was expressed as a fraction of the total lesion area. For morphological analysis of microglia, Iba1+ microglia were identified in *CellProfiler Analyst*, and this population were then further analysed within *CellProfiler* to quantify morphological features of each cell’s Iba1+ mask and skeleton.

To quantify remyelination from toluidine blue-stained resin sections in the probiotic study, slides of the 10 lesions (5 per group) were independently ranked by two experienced, blinded investigators (GG and CZ) according to the extent of remyelination. The assigned numerical order (1-10) was used for subsequent non-parametric statistical tests. To quantify remyelination from electron microscopy images, only axons contained entirely within each field were counted. For these axons, the internal and external diameter of the myelin sheath were then traced using a freehand selection tool in *Fiji*. The g-ratio was calculated as the ratio between the diameters of two circles with areas equal to the internal and external selections respectively. Axons with a circularity <0.7 or aspect ratio >2.5 were excluded from further analysis.

### Statistical analysis

All statistical analysis was carried out using a *Jupyter* Notebook with *Python 2*. *In vivo* experiments contained the following numbers of biological replicates per group: antibiotics lysolecithin study: *n* = 4-6 mice, germ-free cuprizone study: *n* = 4-5 mice, probiotic lysolecithin study: *n* = 3-5 mice. These group sizes were chosen based on previous work and were thought to be sufficiently powered to detect meaningful differences in the OPC / inflammatory response to demyelination. For *in vivo* cell counts, generally 3-4 technical replicate sections were counted and averaged per biological replicate. For *in vitro* cell assays, 3-5 technical replicate wells were averaged for each of 4-5 biological replicate studies.

Data was tested for normality of residuals (Kolmogorov-Smirnov test) and homogeneity of variance (Levene’s test). Data sets passing both of these criteria were compared by either unpaired Student’s *t*-test (if 2 groups), or one-way ANOVA with Tukey HSD *post hoc* tests (if >2 groups). Non-parametric data was compared by Mann-Whitney *U* test (2 groups) or Kruskal-Wallis test with Dunn’s *post hoc* test (>2 groups). For *in vitro* assays, treated conditions were compared to control conditions using a paired-samples *t*-test with the Holm-Bonferroni correction for multiple comparisons. For ranking analysis of remyelination, groups were compared using the Mann-Whitney *U* test. For all statistical tests, differences were considered significant if p<0.05, and the respective test is described in each figure legend.

In all bar plots, the height of the bar represents the group mean, with an error bar representing the standard error of the mean (SEM). *In vivo* data are overlaid with strip plots, in which a grey point represents the value for each individual animal.

## Author contributions

RJMF and DCF conceived the project. CEM, AGF, OBZ, MMH, DCF and RJMF designed the experiments. CEM, AGF, DCF and RJMF wrote and edited the manuscript. CEM designed the figures. CEM, AGF, RP, YD, GAG, CZ and FK conducted the experiments. CEM, AGF, RP, GAG, CZ, FK, AMHY and MMH collected data. CEM and AGF analysed data. CEM, AGF, RP, OBZ, CZ, MMH, DCF and RJMF interpreted data. GAG, CZ, FK and JLG assisted technically. CAJ, CH, DCF and RJMF supervised the project. RJMF and DCF supported this study financially.

## Acknowledgements

We thank Andrew Goldson and Arlaine Brion (Quadram Institute) for their expertise and assistance in running the GF study. We are also grateful to Daniel Morrison, John Falconer and Michal Presz for their technical assistance and to the Cambridge Advanced Imaging Centre for use of their electron microscope. This work was supported by grants from UK Multiple Sclerosis Society, The British Trust for the Myelin Project, MedImmune, The Adelson Medical Research Foundation and a core support grant from the Wellcome Trust and MRC to the Wellcome Trust-Medical Research Council Cambridge Stem Cell Institute. CEM was supported by grants from the Jean Shanks Foundation and the James Baird Fund and OBZ received a BIRAX fellowship.

## Supplementary figures

**Figure 1 Supplement 1:**
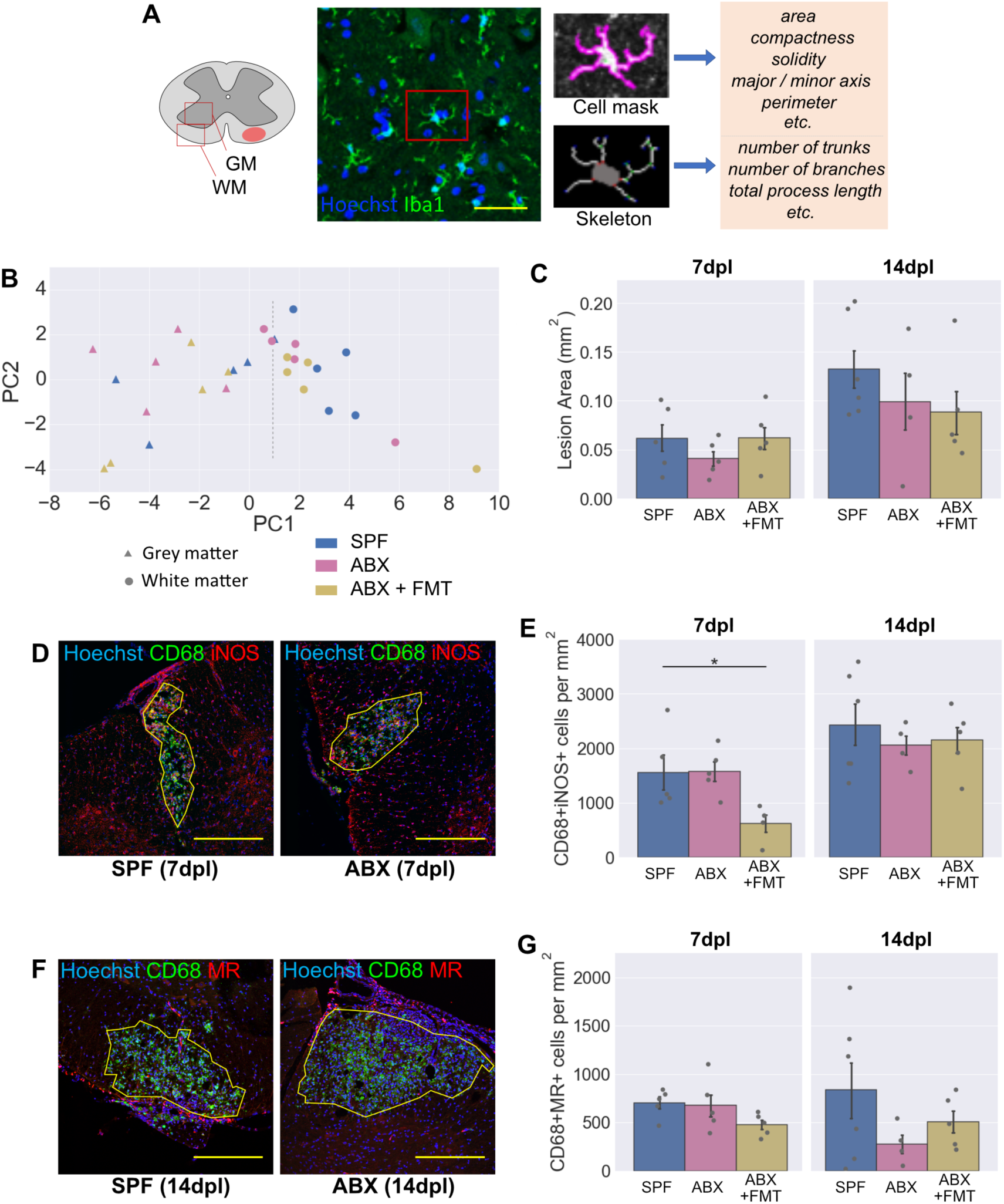
Morphometric analysis of microglia in unlesioned white matter and other features of the inflammatory response to demyelination in ABX-treated mice. (**A**) Protocol for morphometric analysis: spinal cord sections were stained for Iba1, and images taken in the ventral horn grey matter and ventrolateral white matter, contralateral to the lesion. Microglia (Iba1+) were identified using *CellProfiler* software and a number of morphological features were extracted. (**B**) Plot of principal components 1 and 2 from this dataset shows clustering of grey versus white matter microglia (dotted line), but no clear differences between experimental groups. (**C**) Following demyelination, lesion area was not different between groups. (**D-E**) Representative images (**D**) and density (**E**) of iNOS+CD68+ activated microglia / infiltrating macrophages within lesions. (**F-G**) Representative images (**F**) and density (**G**) of MR+CD68+ activated microglia / infiltrating macrophages within lesions. Scale bars: (**A**) = 50μm, (**D**, **F**) = 250μm. Error bars show mean ± SEM; *p<0.05; one-way ANOVA with Tukey HSD *post hoc* test, n=4-6 mice.

**Figure 3 Supplement 1:**
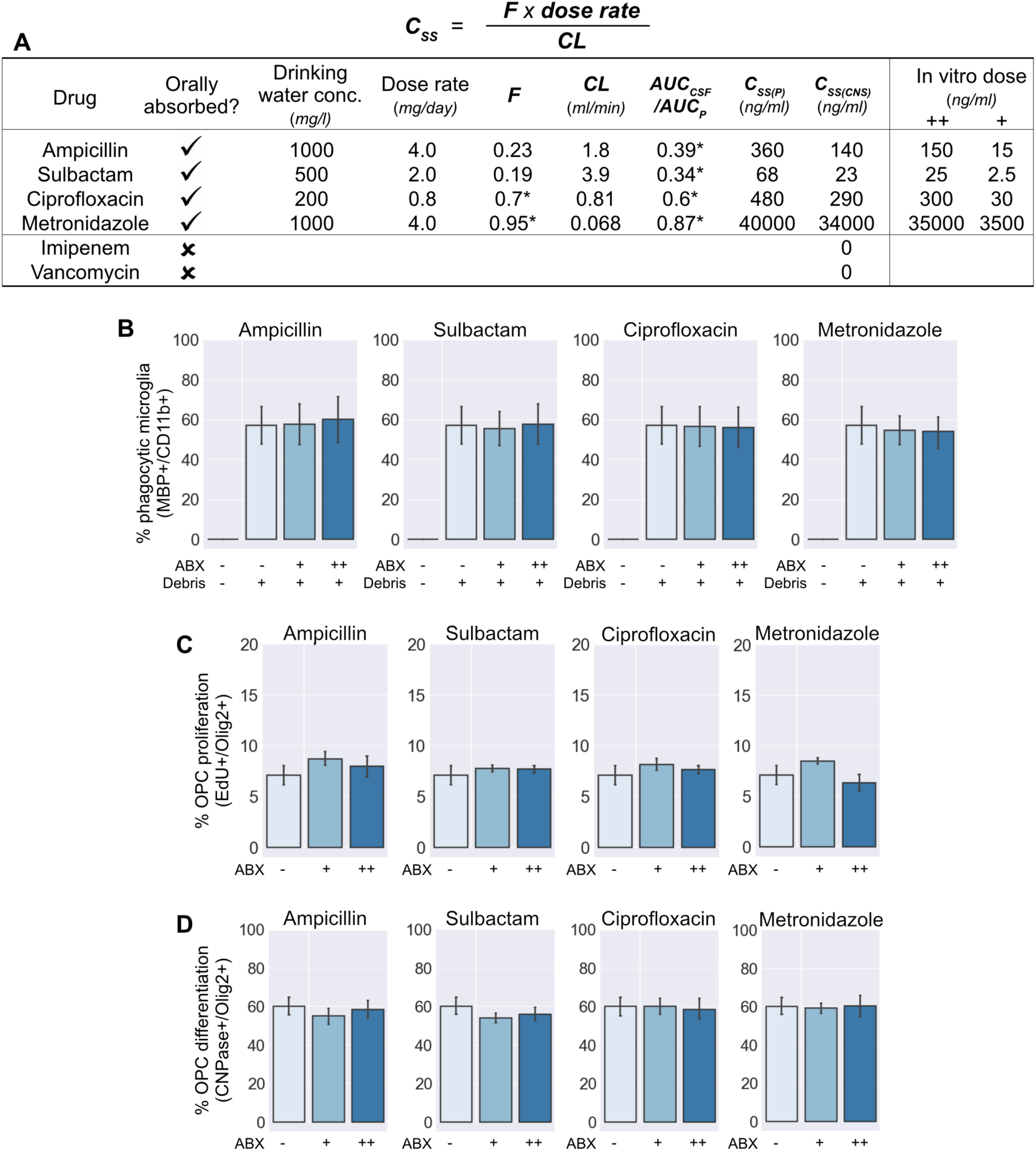
Further data from *in vitro* experiments. (**A**) CNS concentrations (C_SS(CNS)_) of ABX were estimated using literature values for their bioavailability (*F*), clearance (*CL*) and CNS penetration (*AUC*_*CSF*_*/AUC*_*P*_). * denotes values extrapolated from human studies. (**B**) Myelin uptake was quantified for each antibiotic individually at its estimated CNS concentration (++) and a 10% dose (+). (**C-D**) OPC proliferation (**C**) and differentiation (**D**) were quantified for each antibiotic individually at its estimated CNS concentration (++) and a 10% dose (+). Error bars show mean ± SEM; paired-samples *t*-test with Holm-Bonferroni correction n=4-5 separate experiments.

**Figure 4 Supplement 1:**
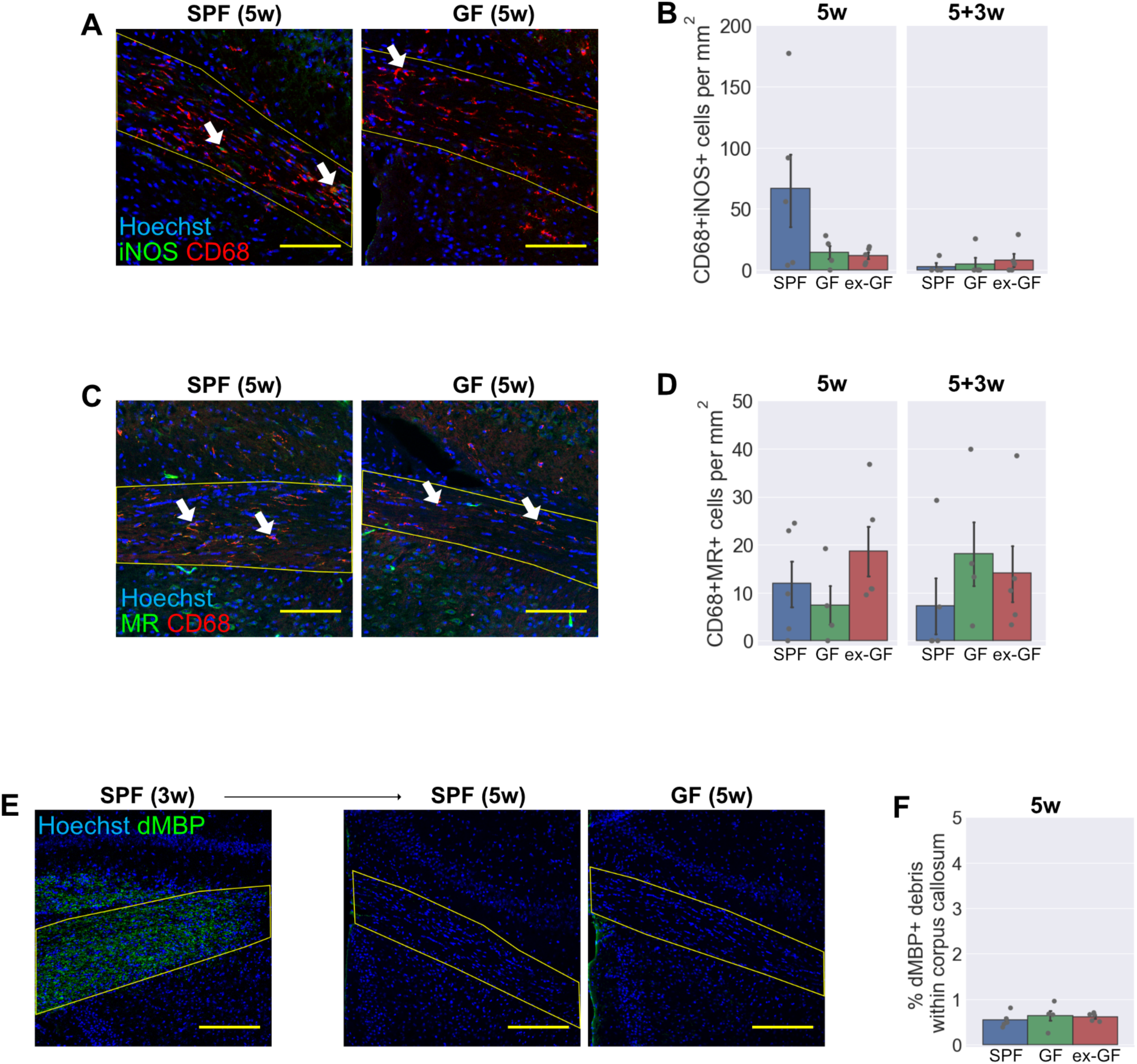
Further data on the inflammatory response to cuprizone in germ-free mice. (**A-B**) Representative images (**A**) and density (**B**) of iNOS+CD68+ activated microglia / infiltrating macrophages within the corpus callosum (yellow line). (**C-D**) Representative images (**C**) and density (**D**) of MR+CD68+ activated microglia / infiltrating macrophages within the corpus callosum. (**E**) After 3 weeks of cuprizone treatment (in a SPF mouse) there was substantial accumulation of dMBP+ myelin debris, which was almost entirely cleared by 5 weeks in both SPF and GF groups. (**F**) Quantification of the remaining dMBP myelin debris after 5 weeks of cuprizone treatment. Scale bars: (**A**, **C**) = 100μm, (**E**) = 200μm. Arrow heads (**A**, **C**) show representative double-positive cells. Error bars show mean ± SEM; one-way ANOVA, n=4-5 mice.

**Figure 5 Supplement 1:**
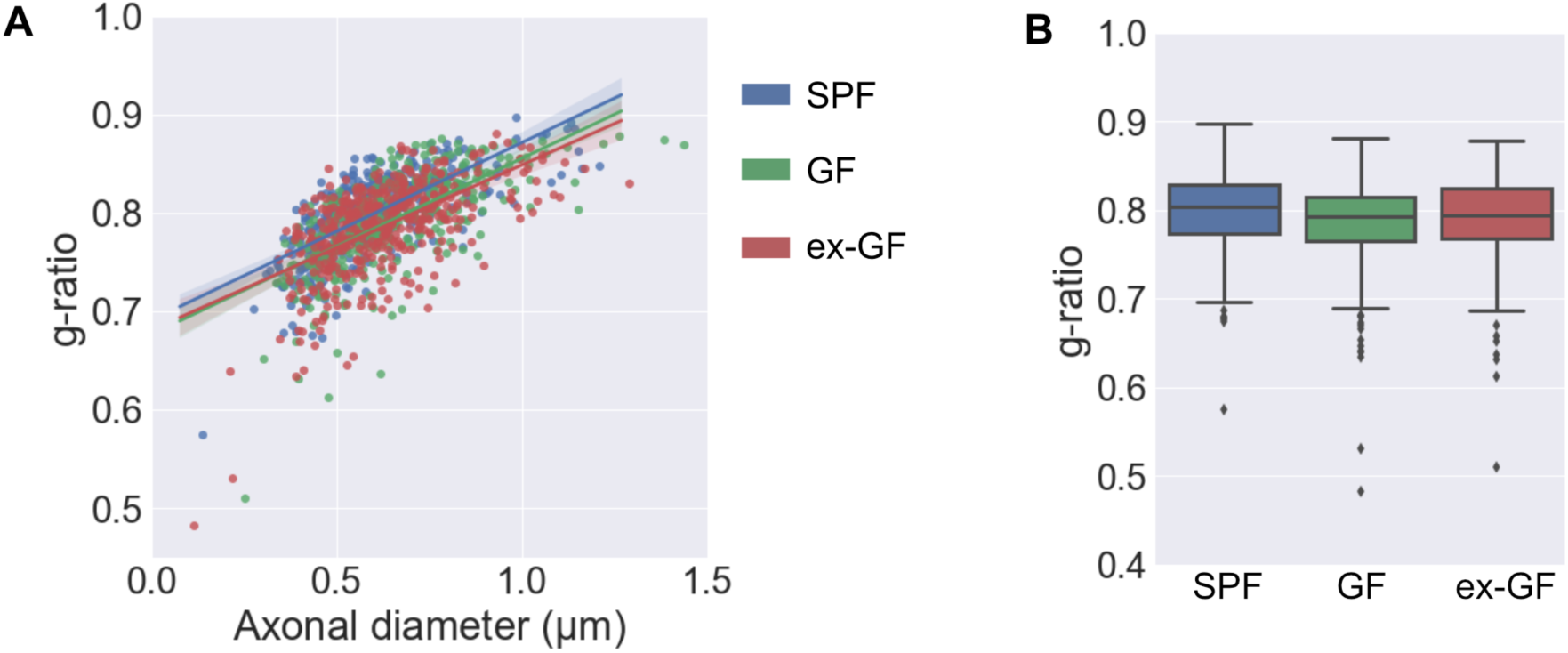
G-ratio analysis of remyelination in germ-free mice. (**A**) g-ratios of all axons analysed, expressed as a function of axonal diameter with a regression line for each group. (**B**) Boxplot of g-ratios separated by experimental group. Shaded areas (**A**) show 95% confidence interval for each line. Outliers (**B**) are more than 1.5 times the interquartile range (IQR) beyond the boundaries of the IQR. SPF n=371 axons; GF n=375 axons; ex-GF n=423 axons.

**Figure 6 Supplement 1:**
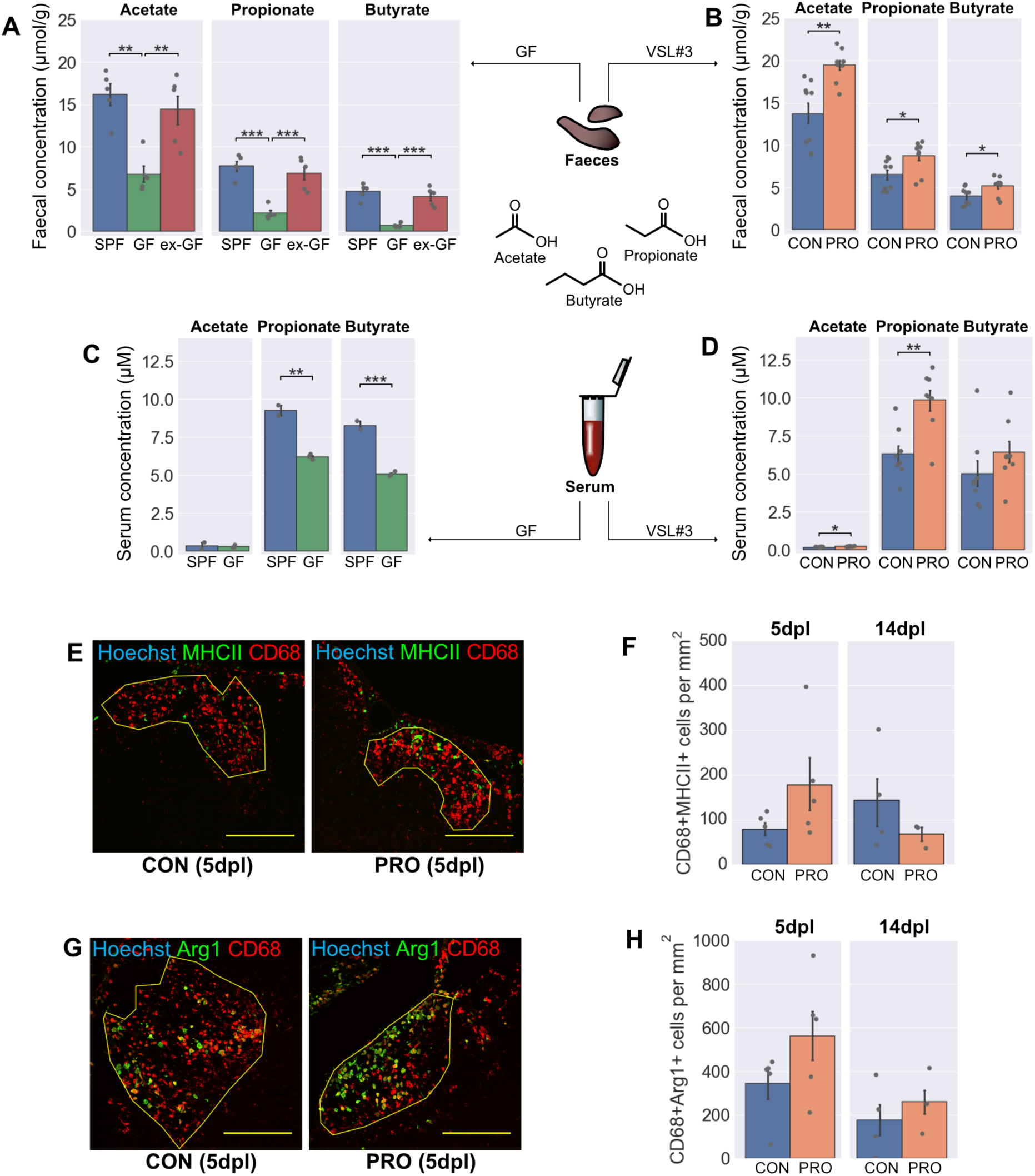
Additional effects of VSL#3 treatment on SCFAs and inflammation. (**A-D**)Quantification by gas chromatography-mass spectroscopy (GC-MS) of SCFAs in the faeces of GF (**A**) and VSL#3-treated mice (**B**), and in the serum of GF (**C**) and VSL#3-treated mice (**D**). (**E-F**) Representative images (**E**) and density (**F**) of MHCII+CD68+ activated microglia / infiltrating macrophages within lesions. (**G-H**) Representative images (**G**) and density (**H**) of Arg1+CD68+ activated microglia / infiltrating macrophages within lesions. Scale bars (**E**, **G**) = 250μm. Error bars show mean ± SEM; *p<0.05, *p<0.01, ***p<0.001; (**A**) one-way ANOVA with Tukey HSD *post hoc* test n=4-5 mice; (**B**, **D**) Student’s *t*-test, n=8 mice; (**C**) Student’s *t*-test, n=2-5 mice; (**F**, **H**) Student’s *t*-test, n=3-5 mice.

**Figure 7 Supplement 1:**
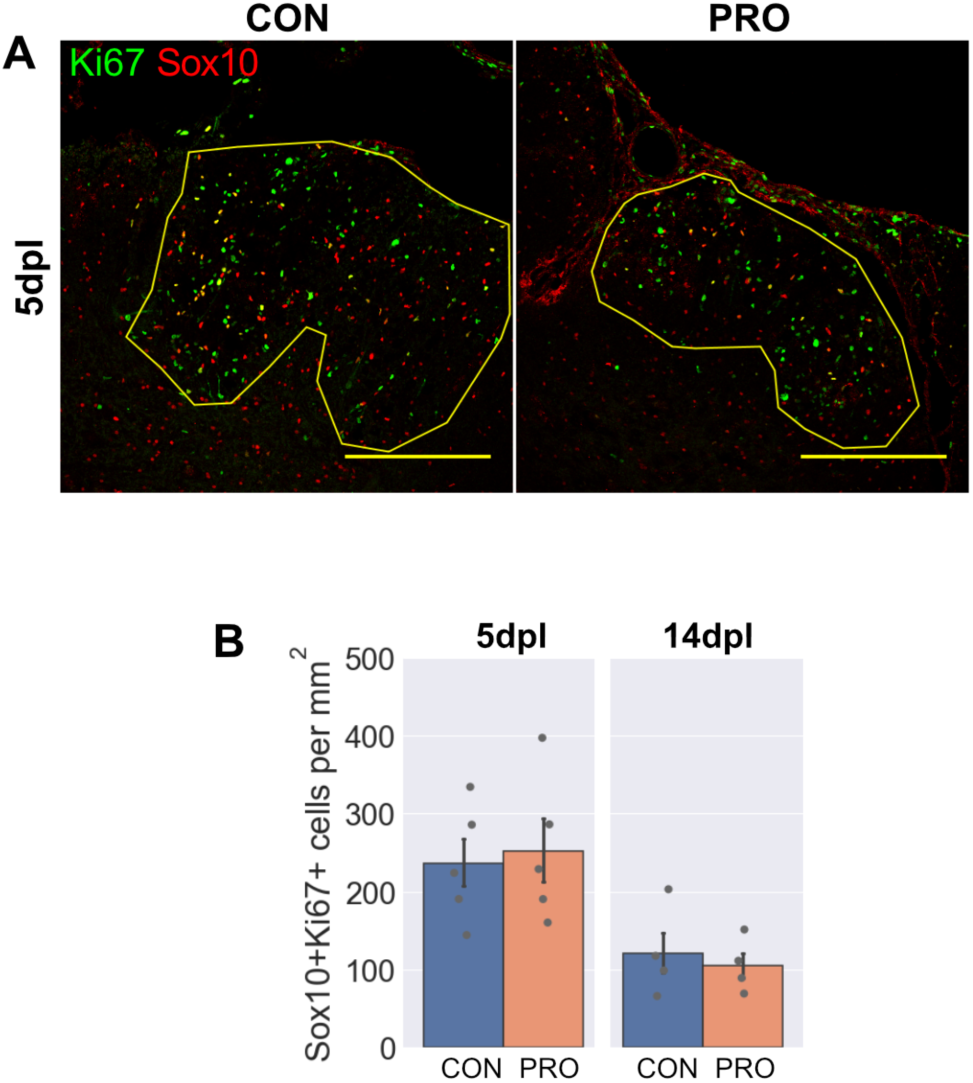
Additional effects of VSL#3 treatment on OPCs. (**A-B**) Representative images (**A**) and density (**B**) of Sox10+Ki67+ proliferating OPCs within lesions. Scale bars (**A**) = 250μm. Error bars show mean ± SEM; n = 3-5 mice.

